# Bioinformatic analysis of brain-specific miRNAs for identification of candidate traumatic brain injury blood biomarkers

**DOI:** 10.1101/794321

**Authors:** Christine Smothers, Chris Winkelman, Grant C. O’Connell

**Affiliations:** Frances Payne Bolton School of Nursing, Case Western Reserve University, Cleveland, OH

## Abstract

**Background:** Detection of brain-specific miRNAs in the peripheral blood could serve as a surrogate marker of traumatic brain injury (TBI). Here, we systematically identified brain-enriched miRNAs, and tested their utility for use as TBI biomarkers in the acute phase of care.

**Methods:** Publically-available microarray data generated from 31 postmortem human tissues was used to rank 1,364 miRNAs in terms of their degree of brain-specific expression. Levels of the top five ranked miRNAs were then prospectively measured in serum samples collected from 10 TBI patients at hospital admission, as well as from 10 controls.

**Results:** The top five miRNAs identified in our analysis (miR-137, miR-219a-5p, miR-128-3p, miR-124-3p, and miR-138-5p) exhibited 31 to 74-fold higher expression in brain relative to other tissues. Furthermore, their levels were elevated in serum from TBI patients compared to controls, and were collectively able to discriminate between groups with 90% sensitivity and 80% specificity. Subsequent informatic pathway analysis revealed that their target transcripts were significantly enriched for components of signaling pathways which are active in peripheral organs such as the heart.

**Conclusions:** The five candidate miRNAs identified in this study have promise as blood biomarkers of TBI, and could also be molecular contributors to systemic physiologic changes commonly observed post-injury.

**A FINAL PEER REVIEWED VERSION OF THIS ARTICLE HAS BEEN PUBLISHED IN *BRAIN INJURY* AT THE FOLLOWING DOI: 10.1080/02699052.2020.1764102**

There are some notable differences between the analysis presented in this preprint and our final peer-reviewed article. There was a single tissue sample originating from spinal cord that we had classified as a non-brain tissue in our original analysis outlined in this preprint. Because the composition of spinal cord and brain are highly similar in terms of gene expression, classifying this sample as a non-brain tissue dramatically reduced the levels of brain enrichment observed in the analysis. Because brain and spinal cord are molecularly highly similar, but technically distinct anatomical structures, we simply decided to exclude this sample from our final analysis published in *Brain Injury* to avoid confounds. The top 5 miRNAs identified in our original analysis still fell within the top 7 of this final analysis. In addition, the final analysis identified two additional miRNAs which could be candidate biomarkers based on levels of brain enrichment.

The final article published in *Brain Injury* also reports an additional confirmatory tissue specificity analysis performed in a second independent dataset, as well as additional analysis examining the brain specificity of several notable previously proposed miRNA TBI biomarkers, which is not described in this preprint.

## Introduction

Traumatic brain injury (TBI) affects an estimated 1.7 million people in the United States each year, and generates an estimated economic burden of $76.5 billion annually in total lifetime direct medical costs and productivity losses [1]. Quick recognition of TBI during triage avoids clinical delay and ensures patients are referred to an appropriate care team which is trained to manage neurological injury; timely and appropriate referral can limit secondary complications and ultimately improve outcome. Unfortunately, the tools available to clinicians for recognition of TBI during early triage are limited. Symptom-based TBI screening tools available to clinicians in the prehospital and early in-hospital setting have been reported to be as low at 30% sensitive, especially in patients with closed head injuries [2,3]. Furthermore, even when these symptom-based tools are combined with CT imaging, mild to moderate TBI cases can be difficult to diagnose [4].

The identification of peripheral blood biomarkers associated with traumatic brain injury (TBI) could lead to the development of molecular diagnostic which could potentially be used to quickly and non-invasively aid in the recognition of TBI during triage. It is becoming increasingly evident that brain tissue exhibits patterns of micro-RNA (miRNA) expression which are distinct from those of other tissues [5]. During TBI, molecules from damaged neural tissues are released into the extracellular environment and my enter peripheral circulation [6]. Thus, the detection of brain-specific miRNAs in peripheral blood could serve as a surrogate marker of TBI.

The use of brain-specific miRNAs as TBI biomarkers has several theoretical advantages. The blood brain barrier inhibits the diffusion of other brain-specific macromolecules such as proteins into the blood [7], limiting their potential as candidate biomarkers for TBI. miRNAs on the other hand, display the ability to cross the blood brain barrier via microvesicles, exosomes, and lipoprotein carriers [8], and thus may be more readily detected in the blood following injury. The detection of brain-specific proteins in the blood may be further confounded by serum protease degradation [9], while miRNAs can exhibit exceptional stability in blood [10]. Furthermore, miRNAs are more amenable to high-throughput profiling then proteins, allowing for more efficient biomarker discovery, and are becoming increasingly close to being testable at the point-of-care [11], which provides a path to clinical implementation.

Despite the potential advantages of using miRNAs as brain-specific biomarkers, much of the prior biomarker discovery in TBI has focused on proteomic analysis, and there have been relatively few miRNA-based TBI investigations in humans, especially in the acute phase of care. Here, we employed a bioinformatic workflow to systematically screen over 1,300 candidate miRNAs to identify those enriched in human brain tissue, and then tested their potential utility for use as acute blood biomarkers of TBI in a small real-world clinical population.

## Materials and methods

### Experimental design

First, we obtained publically-available expression data for 1,364 miRNAs generated from 31 different post-mortem tissues harvested from two human donors. Tissue specificity index was calculated for each miRNA, and they were subsequently ranked in terms of their degree of brain-specific expression. The abundances of the top five ranked miRNAs were then prospectively measured in admission serum samples collected from 10 TBI patients and 10 healthy controls using qPCR, and evaluated for their ability to discriminate between groups. Finally, in order to better understand the potential systemic impact of altered circulating levels of the top five ranked miRNAs, we informatically predicated their target transcripts and determined whether these targets were enriched for components of specific signaling pathways.

### Tissue specificity analysis

Raw microarray data from human tissues generated by Ludwig et al. were directly obtained from https://ccb-web.cs.uni-saarland.de/tissueatlas/. Data were normalized via quantile normalization using the preprocess package for R. miRNAs were filtered to only retain those which exhibited the highest median expression levels in brain relative to other tissues. The remaining miRNAs were then ranked according to the Tau tissue specificity metric described by Yanai et al. [12].

### Patients

Patients were recruited in the emergency department at Ruby Memorial Hospital (Morgantown, WV). TBI patients received a clinical diagnosis of TBI based on clinical evaluation and review of lab and radiographic findings by an experienced neurologist. Severity of TBI was determined by Glasgow Coma Score, as assessed by a trained clinician [13]. Patients under 18 years of age, previously hospitalized within 30 days, or presenting more than 24 hours post-injury were excluded. Control subjects were recruited under similar exclusion criteria as TBI patients, and were free of neurological symptoms. Demographic information was collected from either the subject or significant other by a trained clinician. Procedures were approved by the institutional review boards of West Virginia University and Ruby Memorial Hospital. Written informed consent was obtained from all subjects or their authorized representatives prior to study procedures.

### Blood sampling

Peripheral venous blood samples were collected from subjects via gel barrier serum separator vacutainers (Becton Dickenson, Franklin Lakes, NJ). Blood was allowed to clot for a minimum of 30 minutes, and vacutainers were spun at 2,000*g for 10 minutes at room temperature. The resultant serum was aliquoted and immediately stored at -80C until analysis.

### miRNA extraction and qPCR

300 uL of serum was spiked with *C. elegans* miR-39, and miRNA was isolated via trizol extraction and subsequent spin column clean-up (miRNeasy, Qiagen, Germantown, MD). Reverse transcription was performed via the miScript II reverse transcription kit (Qiagen). qPCR was performed using the RotorGeneQ thermocycling system (Qiagen) using the miScript SYBR green master mix (Qiagen) and commercially available miScript primer assays (Qiagen) targeting miR-39 and the 5 top ranked candidate miRNAs. CT values of candidate miRNAs were normalized via those of the miR-39 spike-in control. miRNA levels are reported as absolute fold difference relative to the mean of the control group.

### miRNA target prediction and pathway enrichment analysis

The target transcripts of the top ranked miRNAs were predicted using TargetScan [14], and STRING [15] was used query both KEGG [16] and Reactome [17] pathway annotations to determine whether target transcripts were enriched for those involved in specific signaling pathways.

### Statistical analysis

All statistics were performed using R 3.4 [18]. Fisher’s exact test was used for comparison of dichotomous variables, while t-test was used for the comparison of continuous variables. The combined ability of the serum levels of candidate miRNAs to discriminate between stroke patients and controls was tested using k-nearest neighbors (k-NN). Classification was performed using z-scored expression values, three nearest neighbors, and majority rule via the knn.cv() function of the “class” package. The resultant prediction probabilities were used to generate receiver operator characteristic (ROC) curves using the roc() function of the “pROC” package [19]. In the case of all statistical testing, the null hypothesis was rejected when p<0.05. The parameters of all statistical tests performed are outlined in detail within the figure legends.

## Results

### Tissue specificity analysis

Figure 1 depicts the expression levels of the top 50 most brain-enriched miRNAs in each of the 31 tissue types interrogated in our analysis. The top five miRNAs, miR-137, miR-219a-5p, miR-128-3p, miR-124-3p, and miR-138-5p, exhibited 31 to 74-fold higher expression in brain relative to other tissues.

**Figure 1.**
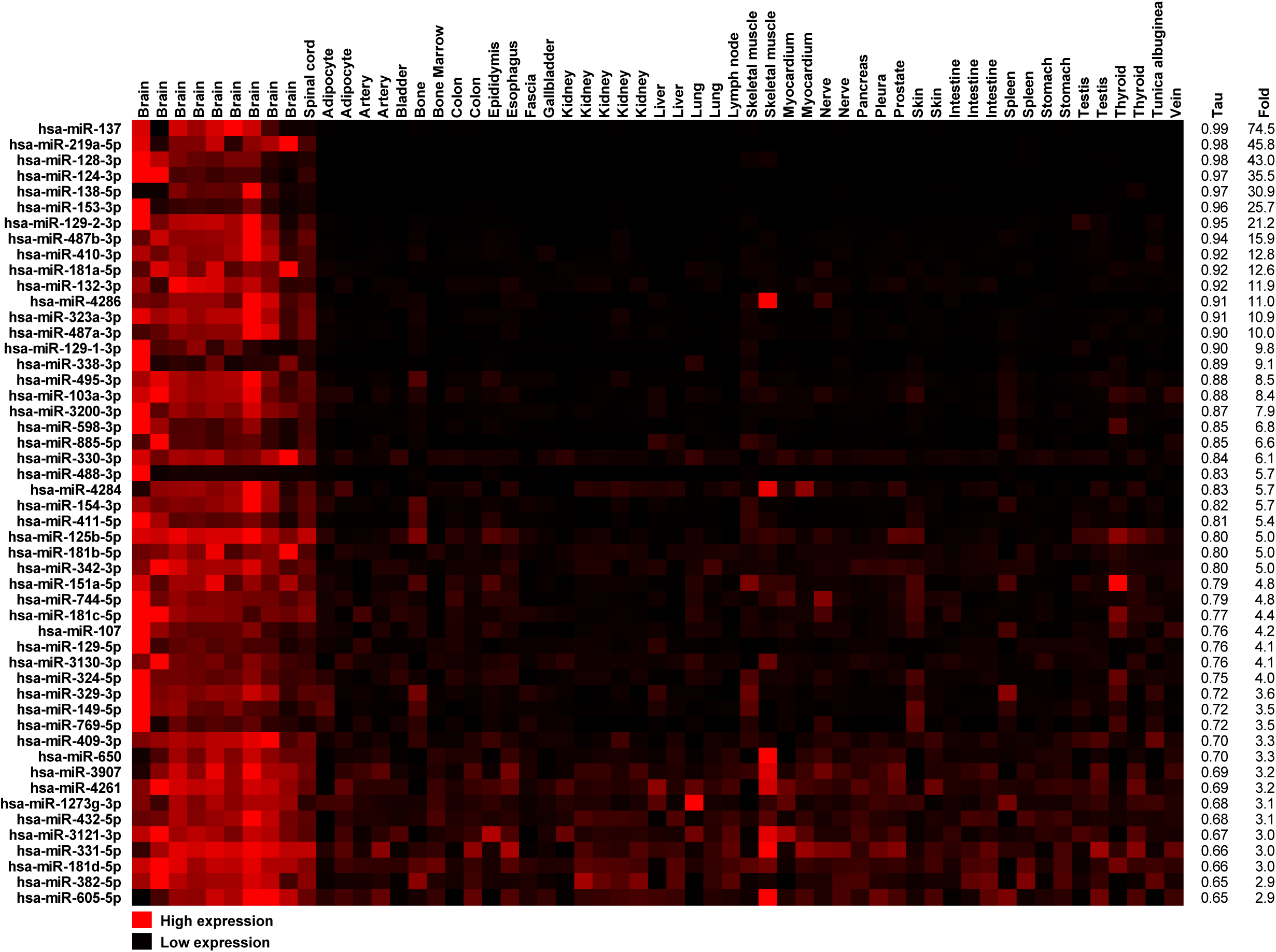
miRNA expression levels across human tissues. Expression levels of the top 50 most brain-enriched miRNAs across samples from 31 distinct human tissues, along with tissue specificity index (Tau) and fold enrichment in brain samples relative to samples from other tissues.

### Diagnostic performance of candidate miRNAs

Clinical and demographic characteristics of TBI and control patients are depicted in Table 1. TBI and control patients were highly similar in terms of similar age and ethnicity. While not statistically significant, the TBI group did contain a higher proportion of female subjects, and a higher rate of hypertension, compared to the control group.

**Table 1.** Clinical and demographic characteristics.

|  | **Control (n=10)** | **TBI (n=10)** | **p-value** |
| --- | --- | --- | --- |
| **Age<sup>a</sup> mean ± SD** | 50.9 ± 16.4 | 50.9 ± 23.7 | 1 |
| **Female<sup>b</sup> n (%)** | 4 (40) | 7 (70) | 0.369 |
| **Caucasian<sup>b</sup> n (%)** | 10 (100) | 10 (100) | 1 |
| **Hypertension<sup>b</sup> n (%)** | 0 (0) | 4 (40) | 0.087 |
| **Dyslipidemia<sup>b</sup> n (%)** | 0 (0) | 1 (10) | 1 |
| **Diabetes<sup>b</sup> n (%)** | 1 (10) | 2 (20) | 1 |
| **Smoking<sup>b</sup> n (%)** | 2 (20) | 4 (40) | 0.629 |

Intergroup comparison via <sup>a</sup>two-sample two-tailed t-test or <sup>b</sup>Fisher's exact test

All five candidate miRNAs exhibited higher levels in serum from TBI patients relative to serum from control patients, however, only three of these differences were statistically significant (Figure 2). The combined serum levels of the five candidate miRNAs were able to discriminate between TBI patients and control patients with 90% sensitivity (0.95 CI: 75.8-100%) and 80% specificity (0.95 CI: 65.8-95.3%) using k-NN (Figure 3).

**Figure 2.**
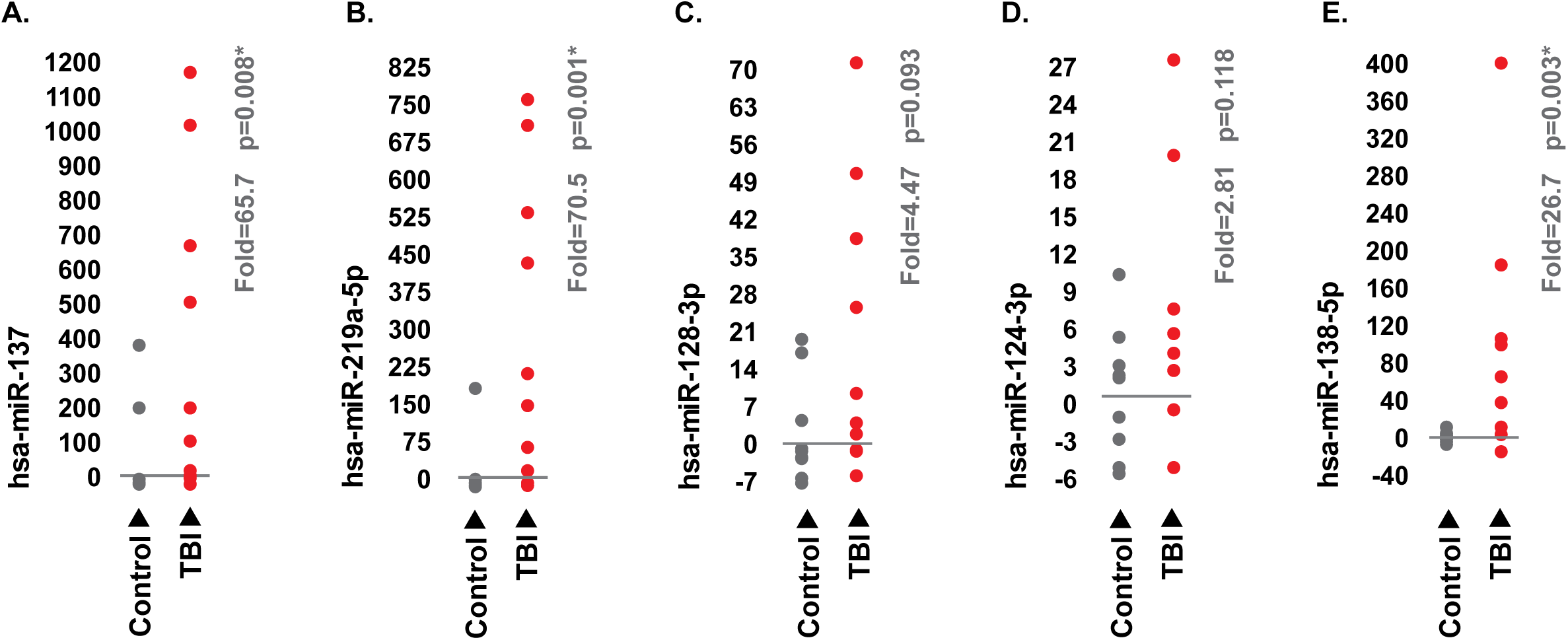
Expression levels of top 5 candidate miRNAs in TBI patients and controls. (A-G) Levels of miR-137, miR-219a-5p, miR-128-3p, miR-124-3p, and miR-138-5p in serum samples from TBI patients and controls collected at emergency department admission. miRNA levels are reported as absolute fold difference relative to the mean of the control group. Means were compared between groups using two-sample two-tailed t-test.

**Figure 3.**
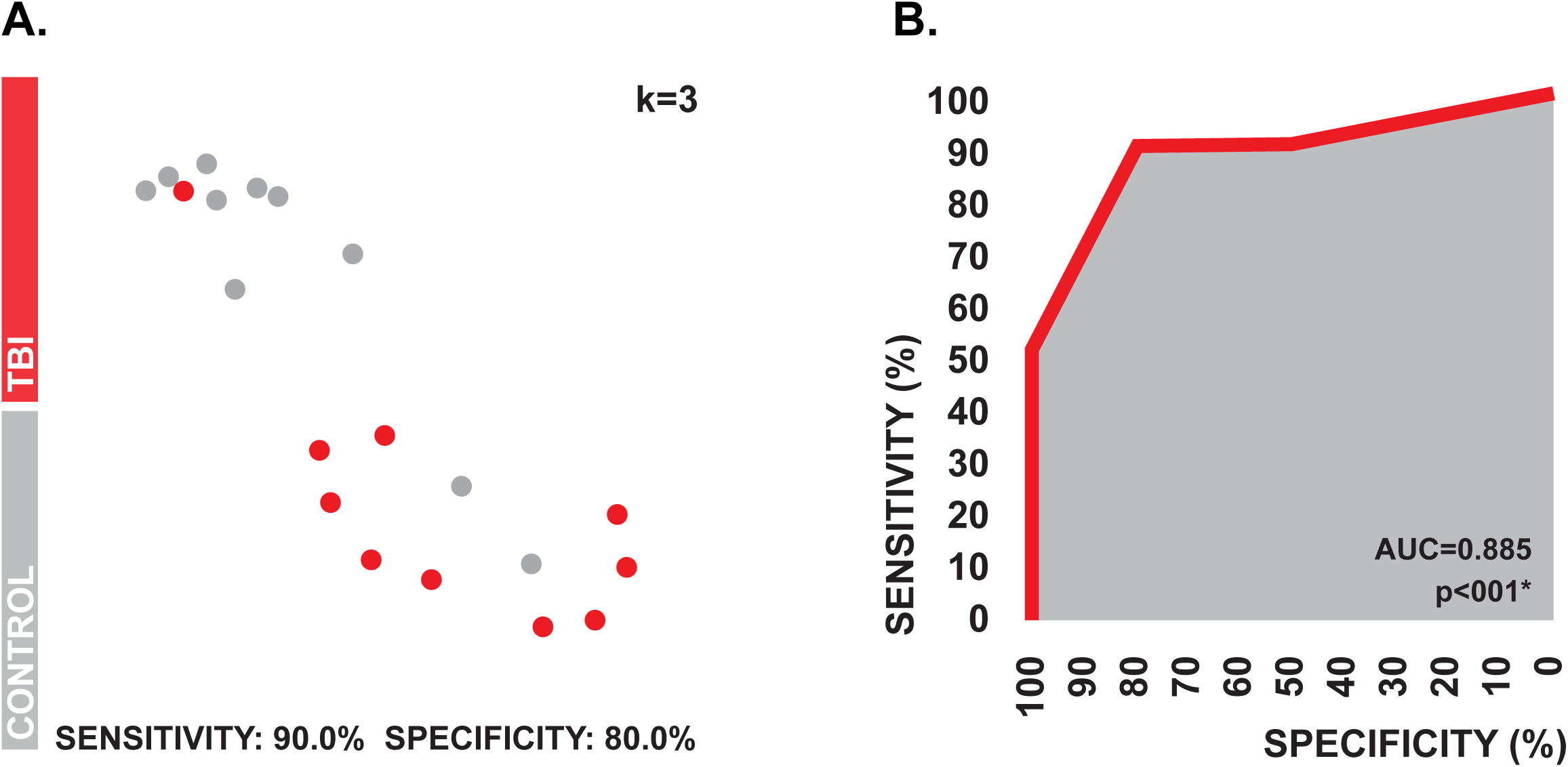
Diagnostic performance of the top 5 candidate miRNAs. (A) Two-dimensional projection of the kNN feature space generated by the coordinate serum levels of the 5 candidate miRNAs. Levels of sensitivity and specificity are associated with class predications generated via three nearest neighbors using a probability cutoff of 0.50. (B) ROC curve associated with the prediction probabilities generated in kNN.

### Biologic function of candidate miRNAs

The predicted targets of the top 5 miRNAs were significantly enriched for transcripts involved signaling pathways which control a wide range of biologic processes such as cellular proliferation and differentiation, neuronal signal transduction, and inflammation (Table 2). Unsurprisingly, several of these signaling pathways were specific to CNS processes, however, they also included those which are active in non-CNS tissues. For example, the predicted targets of miR-137 were significantly enriched for transcripts which are involved in cardiac-specific signaling pathways (Figure 4). Broadly, this suggests that miRNAs released from the brain into circulation during TBI could alter molecular signaling in peripheral organs, which could ultimately help drive some of the distal physiologic changes which are commonly observed post-injury such as cardiac dysfunction.

**Table 2.** Signaling pathways targeted by the top five candidate miRNAs.

| miRNA: | Number of target mRNAs<sup>a</sup>: | Enriched pathways<sup>b</sup>: | Database: | Accession number: | p-value<sup>c</sup>: | Neural-specific pathway: |
| --- | --- | --- | --- | --- | --- | --- |
| **miR-137** | 1,314 | ErbB signaling pathway | KEGG | hsa04012 | 0.0046 | No |
|  |  | Axon guidance | KEGG | hsa04360 | 0.0046 | Yes |
|  |  | RAF/MAP kinase cascade | Reactome | HSA-5673001 | 0.0193 | No |
|  |  | MAPK signaling pathway | KEGG | hsa04010 | 0.0208 | No |
|  |  | Calcium signaling pathway | KEGG | hsa04020 | 0.0208 | No |
|  |  | cAMP signaling pathway | KEGG | hsa04024 | 0.0208 | No |
|  |  | Thyroid hormone signaling pathway | KEGG | hsa04919 | 0.032 | No |
|  |  | Cardiac conduction | Reactome | HSA-5576891 | 0.0454 | No |
| **miR-219a-5p** | 449 | Hippo signaling pathway | KEGG | hsa04390 | 0.0172 | No |
| **miR-128-3p** | 1,225 | Pluripotency of stem cells | KEGG | hsa04550 | 0.00089 | No |
|  |  | Axon guidance | KEGG | hsa04360 | 0.0024 | Yes |
|  |  | MAPK signaling pathway | KEGG | hsa04010 | 0.0051 | No |
|  |  | Rap1 signaling pathway | KEGG | hsa04015 | 0.0051 | No |
|  |  | Ras signaling pathway | KEGG | hsa04014 | 0.0083 | No |
|  |  | FoxO signaling pathway | KEGG | hsa04068 | 0.0107 | No |
|  |  | TGF-beta signaling pathway | KEGG | hsa04350 | 0.0107 | No |
|  |  | Thyroid hormone signaling pathway | KEGG | hsa04919 | 0.0118 | No |
|  |  | Neurotrophin signaling pathway | KEGG | hsa04722 | 0.0122 | Yes |
|  |  | AMPK signaling pathway | KEGG | hsa04152 | 0.0157 | No |
|  |  | Phospholipase D signaling pathway | KEGG | hsa04072 | 0.019 | No |
|  |  | Insulin signaling pathway | KEGG | hsa04910 | 0.019 | No |
|  |  | Apelin signaling pathway | KEGG | hsa04371 | 0.0305 | No |
|  |  | ErbB signaling pathway | KEGG | hsa04012 | 0.0351 | No |
|  |  | Circadian entrainment | KEGG | hsa04713 | 0.0351 | Yes |
|  |  | cAMP signaling pathway | KEGG | hsa04024 | 0.0381 | No |
|  |  | TNF signaling pathway | KEGG | hsa04668 | 0.043 | No |
|  |  | Neurexins and neuroligins | Reactome | HSA-6794361 | 0.0442 | Yes |
| **miR-124-3p** | 1,820 | Axon guidance | KEGG | hsa04360 | 0.0021 | Yes |
|  |  | Neurotrophin signaling pathway | KEGG | hsa04722 | 0.0021 | Yes |
|  |  | Signaling by NTRKs | Reactome | HSA-166520 | 0.004 | No |
|  |  | Sphingolipid signaling pathway | KEGG | hsa04071 | 0.0045 | No |
|  |  | Cellular senescence | KEGG | hsa04218 | 0.0065 | No |
|  |  | Apelin signaling pathway | KEGG | hsa04371 | 0.0084 | No |
|  |  | Insulin signaling pathway | KEGG | hsa04910 | 0.0084 | No |
|  |  | ErbB signaling pathway | KEGG | hsa04012 | 0.0099 | No |
|  |  | Regulation of actin cytoskeleton | KEGG | hsa04810 | 0.0277 | No |
|  |  | Oxytocin signaling pathway | KEGG | hsa04921 | 0.0277 | Yes |
|  |  | Pluripotency of stem cells | KEGG | hsa04550 | 0.0342 | No |
|  |  | cGMP-PKG signaling pathway | KEGG | hsa04022 | 0.0493 | No |
| **miR-138-5p** | 704 | NOTCH signaling | Reactome | HSA-157118 | 0.0073 | No |
|  |  | Cellular senescence | Reactome | HSA-2559583 | 0.0097 | No |
|  |  | Axon guidance | Reactome | HSA-422475 | 0.0241 | Yes |
|  |  | Circadian clock | Reactome | HSA-400253 | 0.0248 | Yes |
|  |  | Netrin-1 signaling | Reactome | HSA-373752 | 0.0476 | No |

<sup>a</sup>A complete list of predicted target transcripts can be found in Supplemental File 1; <sup>b</sup>Only select pathways are shown, a complete list of enriched pathways can be found in Supplemental File 2; <sup>c</sup>P-values adjusted for multiple comparisons using the Benjamini-Hochberg method to control false discovery rate.

**Figure 4.**
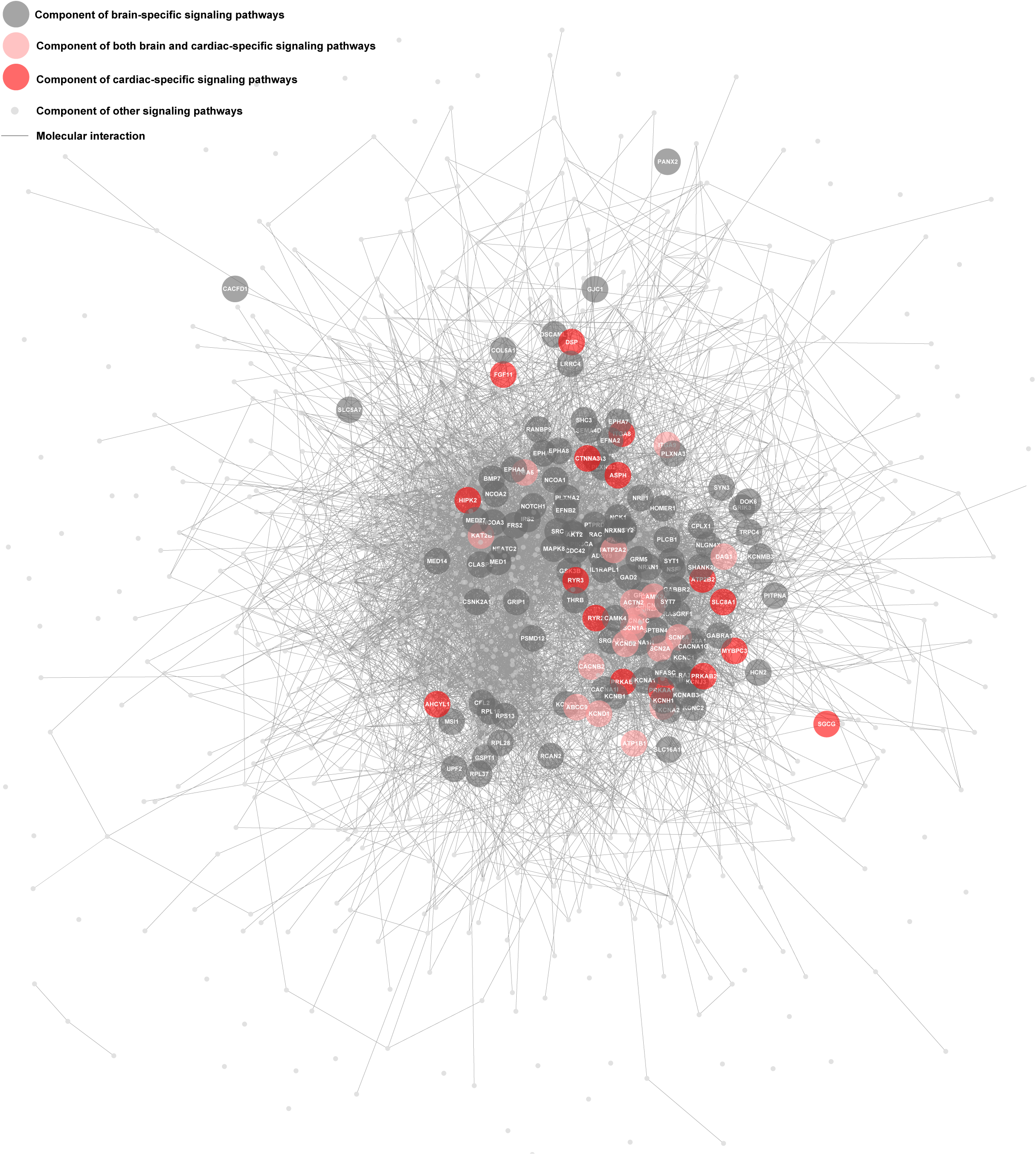
Components of brain-specific and cardiac specific signaling pathways targeted by miR-137. Molecular interaction network depicting the relationships between the protein products of transcripts targeted by miR-137. Components of significantly enriched brain-specific and cardiac-specific signaling pathways are indicated.

## Discussion

Blood biomarkers associated with traumatic brain injury (TBI) could be used to non-invasively aid in the recognition of TBI during triage. The detection of brain-specific miRNAs released from damaged neural tissue into the peripheral blood could serve as a surrogate marker of TBI. Here, we employed a systematic analysis identify brain-enriched miRNAs, and then tested their potential utility for use as TBI blood biomarkers in the acute phase of care. Our analysis identified several candidate miRNAs that have yet to be explored as TBI biomarkers, and the levels of diagnostic performance which we observed suggest that they could have future clinical utility. Furthermore, our informatics analysis suggests that circulating miRNAs originating from damaged brain tissue could be molecular contributors to the peripheral physiologic changes which are commonly observed in TBI.

To our knowledge, the five candidate miRNAs identified in our analysis have yet to be widely investigated as biomarkers in the acute phase of human TBI. We observed elevated serum levels of all five candidate miRNAs (miR-137, miR-219a-5p, miR-128-3p, miR-124-3p, and miR-138-5p) at hospital admission. These results are constant with a recent study by Yan et al. which reported a similar elevation in circulating levels of miR-219a-5p in human TBI within 24 hours of injury [20], as well as with prior investigations reporting altered circulating levels of miR-137 and miR-124-3p in animal models of TBI [21,22].

The circulating levels of the top 5 miRNAs identified in our analysis were able to differentiate between TBI patients and controls with 90% sensitivity and 80% specificity in the acute phase of care. S100 calcium-binding protein B (S100B) and glial fibrillary acidic protein (GFAP), both proteins which exhibit enriched expression in neural tissue, are two of the most widely investigated protein TBI blood biomarkers; pooled diagnostic accuracy data across multiple clinical investigations suggests that S100B is roughly 85% sensitive and 60% specific for TBI, while GFAP is roughly 80% sensitive and 80% specific [23]. Furthermore, the symptom-based TBI screening tools available to clinicians during early triage have been reported to be roughly 30-45% sensitive and 70-90% specific [2,3]. Albiet in limited sample size, the five candidate miRNAs identified in our analysis displayed levels of diagnostic accuracy which compare favorably to those reported for other candidate TBI biomarkers, as well as symptom-based recognition scales, suggesting that they could have true clinical utility if our findings can be validated in larger patient population.

The experimental strategy employed in this investigation has several distinct advantages over those employed in the small number of large-scale miRNA biomarker discovery analyses performed in TBI to date. Most notably, prior investigations have identified candidate miRNAs by high-throughput comparisons of circulating miRNAs between TBI patients and healthy controls [20,24–26]. Thus, it is unclear whether the miRNAs identified in these studies exhibit elevation in blood which are specific to brain injury, or just trauma in general. Because the five candidate miRNAs which we identified in our analysis exhibit highly localized expression in brain tissue, it much more likely that their presence in blood is directly associated with neurological damage. Furthermore, the use of microarray for discovery screening of miRNAs directly in blood in prior studies may have limited their ability to detect miRNAs originating from brain [24,26]. Brain-specific miRNAs are likely present in circulation at relatively low levels, and microarray is less sensitive than PCR-based methods [27]; thus, differences in circulating levels of brain-specific miRNAs may have been missed due to high lower limits of detection. Here we took advantage to the throughput afforded by microarray to screen miRNAs directly in brain tissue where brain-specific species are much more abundant, and then used PCR for more sensitive detection when analyzing blood samples.

While our findings our findings are promising, it is important to note that this study is not without limitations. Most notably is the small sample size employed in our validation analysis, and the fact our control group was comprised of healthy subjects. In order to determine the true diagnostic accuracy of the five candidate miRNAs identified in our analysis, they need to be investigated further in a larger patient population, ideally including more mild TBI cases and a control group which includes non-neurological trauma patients. Furthermore, we only examined the candidate markers in the acute phase of care and we did not investigate clinical outcomes outside of diagnosis. Future work should examine these miRNAs longitudinally post-injury and explore associations with other clinical measures to determine whether they could be useful in guiding other care decisions. For example, the miRNAs identified in this study might have further utility to quantify axonal injury, monitor secondary tissue damage, or predict adverse complications such increased intracranial pressure.

From a biological perspective, our results may provide an avenue for future study aimed at better understanding the molecular mechanisms which drive the systemic response to TBI. TBI triggers body-wide changes in metabolism [28], immune function [29], and cardiovascular dynamics [30], all of which influence clinical outcome. While the top 5 candidate miRNAs described here exhibit localized expression in brain, our pathway enrichment analysis suggests that they have the ability to modulate several signaling pathways which are active in non-neural peripheral tissues. Due to the ability of circulating miRNAs to enter cells [31], it is possible that the elevations in serum levels of the candidate miRNAs which we observed in TBI could alter gene expression levels in circulating cell populations and peripheral tissues. Better understanding the peripheral consequences that TBI-induced elevations in circulating levels of these miRNAs may have could help identify new therapeutic targets which could be used to modulate the systemic injury response to improve prognosis. For example, the predicted targets of miR-137 were significantly enriched for transcripts involved in signaling pathways which mediate cardiac conduction; thus, it is possible that TBI-induced elevations in circulating miR-137 could play a role in the pathogenesis of post-TBI cardiac dysfunction, a common and serious complication of TBI [32].

Taken as a whole, our results suggest that the five candidate miRNAs identified in our analysis are have promise for use as TBI biomarkers. They displayed levels of diagnostic accuracy which compare favorably to those achievable via previously described candidate TBI biomarkers, as well as symptom-based TBI recognition tools, suggesting that they could have true utility for triage if our preliminary findings can be validated. Furthermore, future exploration into the systemic effects of TBI-induced elevations of these miRNAs could help better understand the peripheral physiologic changes that occur in response to injury.

## Acknowledgements

The authors would like to thank Dr. Paul D. Chantler of West Virginia University, and Dr. Taura L. Barr, formally of West Virginia University, for providing access to blood samples.

## Data availability

Human brain miRNA expression data are available at: https://ccb-web.cs.uni-saarland.de/tissueatlas/. The remaining data that support the findings of this study are available on request from the corresponding author.

## Supplemental Files

*Supplemental File 1:*

https://drive.google.com/open?id=12VVWCpmvw2-_QKr-F9hWMaBCuEVYwmTQ

*Supplemental File 2:*

https://drive.google.com/open?id=1c5En3Wb9b-0An2-5yd6i377RxeRDMnpi

